# Measuring MEG closer to the brain: Performance of on-scalp sensor arrays

**DOI:** 10.1101/073585

**Authors:** Joonas Iivanainen, Matti Stenroos, Lauri Parkkonen

## Abstract

Optically-pumped magnetometers (OPMs) have recently reached sensitivity levels required for magnetoencephalography (MEG). OPMs do not need cryogenics and can thus be placed within millimetres from the scalp into an array that adapts to the invidual head size and shape, thereby reducing the distance from cortical sources to the sensors. Here, we quantified the improvement in recording MEG with hypothetical on-scalp OPM arrays compared to a 306-channel state-of-the-art SQUID array (102 magnetometers and 204 planar gradiometers).

We simulated OPM arrays that measured either normal (nOPM; 102 sensors), tangential (tOPM; 204 sensors), or all components (aOPM; 306 sensors) of the magnetic field. We built forward models based on magnetic resonance images of 10 adult heads; we employed a three-compartment boundary element model and distributed current dipoles evenly across the cortical mantle.

Compared to the SQUID magnetometers, nOPM and tOPM yielded 7.5 and 5.3 times higher signal power, while the correlations between the field patterns of source dipoles were reduced by factors of 2.8 and 3.6, respectively. Values of the field-pattern correlations were similar across nOPM, tOPM and SQUID gradiometers. Volume currents reduced the signals of primary currents on average by 10%, 72% and 15% in nOPM, tOPM and SQUID magnetometers, respectively. The information capacities of the OPM arrays were clearly higher than that of the SQUID array. The dipole-localization accuracies of the arrays were similar while the minimum-norm-based point-spread functions were on average 2.4 and 2.5 times more spread for the SQUID array compared to nOPM and tOPM arrays, respectively.

## 1. Introduction

Magnetoencephalography (MEG) is a non-invasive neuroimaging technique that detects the magnetic fields of electrically active neuron populations in the human brain (Hämäläinen et al., 1993). Due to the weakness of these fields, highly sensitive detectors (magnetometers) are needed. Until recently, superconducting quantum interference devices (SQUIDs) have been the only sensors with adequate sensitivity to enable practical mapping of the cerebral neuromagnetic fields. However, SQUIDs require a cryogenic environment, and liquid helium (boiling point 4.2 K) is typically used for reaching sufficiently low temperatures. The necessity of cryogenics imposes several problems: First, cryogenics make MEG systems bulky. Second, the necessary thermal isolation between the sensors and the subject sets a lower limit, typically about 2 cm, on the distance from the sensors to the scalp of the subject. Last, a SQUID-based sensor array is not adjustable to individual head size and shape, which further increases the average distance between sensors and scalp. Alternative sensors such as optically-pumped, or atomic, magnetometers (OPMs) (Budker and Kimball, 2013; Budker and Romalis, 2007) and high-T_c_ SQUIDs (Öisjöen et al., 2012) have recently reached sensitivities required for detecting neuromagnetic fields.

Particularly OPMs operating in the spin-exchange relaxation-free (SERF) regime (Allred et al., 2002) could become a feasible, non-cryogenic alternative to SQUIDs for neuromagnetic measurements as the combined dimensions and sensitivies of the SERF-OPMs are approaching those of SQUIDs (~4-6 cm^2^ and ~3-5 fT/ 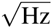). Johnson and colleagues (2010) reported that they achieved a sensitivity better than 5 fT/ 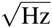 with their SERF-OPM with a footprint of 6×6 cm^2^. A smaller chip-scale microfabricated OPM developed by Mhaskar and colleagues (2012) has a volume of about (1 cm)^3^ and sensitivity better than 20 fT/ 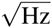. Shah and Wakai (2013) have demonstrated an OPM with a volume of 2×2×5 cm^3^ and sensitivity of 10 fT/ 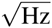 during a field study and 6 fT/ 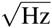 in the laboratory; the theoretical photon-shot-noise-limited sensitivity of that OPM was 1 fT/ 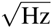.

The sensitivies of the OPMs should thus allow the measurement of weak neuromagnetic fields while their dimensions enable multichannel whole-head-covering measurements. OPMs could also be placed within few millimetres from scalp, permitting EEG-cap-like MEG arrays. Certain aspects of performance of such on-scalp MEG sensor arrays has been recently evaluated by Boto and colleagues (2016); they showed that multichannel whole-head OPM system that measures the magnetic field component normal to the local scalp surface offers a fivefold improvement in sensitivity and clear improvements in reconstruction accuracy and spatial resolution of beamforming compared to traditional SQUID arrays. Our simulations go beyond that work by evaluating also OPM arrays that measure tangential field components, by investigating how the different aspects of MEG signal generation are reflected in the field components and by introducing measures that can be used to quantify and benchmark performance of MEG arrays and which also serve as means to understand the performance differences between arrays. We base our resolution analysis both on a physical model of the measurement set-up and on minimum-norm-based point-spread functions.

We simulate the performance of hypothetical on-scalp OPM arrays of sensors that measure the normal and tangential components of the neuromagnetic field. We compare the performance of such arrays to a state-of-the-art SQUID-based 306-channel MEG array. In our simulations, we use realistic geometries derived from magnetic resonance images of ten adult subjects and calculate magnetic fields using the boundary element method. We use both novel and well-established metrics derived from forward models and point-spread functions of the minimum-norm estimates. The metrics quantify signal power, information, similarity of the field patterns of the sources, overlap of the sensor lead fields, localization accuracy and resolution. Besides measures of performance evaluation, such metrics can be used to guide the design of MEG sensor arrays. In addition, we quantify the differences in the measurements of normal and tangential field components by analyzing the contributions they receive from primary and secondary currents.

## 2. Theory

In this section, we shortly review the key physics and introduce the concepts used in the forward and inverse modeling employed in this study.

### 2.1 Physics

When bioelectromagnetic fields are modeled in the macroscopic scale, the time-dependent terms in Maxwell’s equations can be omitted (Plonsey and Heppner, 1967). The total current density in the head is divided into two parts; **J** = **J**_p_ + **J**_V_, where **J**_p_ is the primary current density representing the neuronal source activity while J_V_ is the volume current density, which is driven by the electric field caused by **J**_p_ and is present everywhere in the conductor. The electric potential φ is then given by ∇·(σ∇φ)=∇·**J**_p_, where σ is conductivity, and the magnetic field can be obtained by integrating the total current using Biot-Savart law. When the conducting region is modeled to consist of piecewise homogeneous regions, the magnetic field is given by the Geselowitz formula; (Geselowitz, 1970)

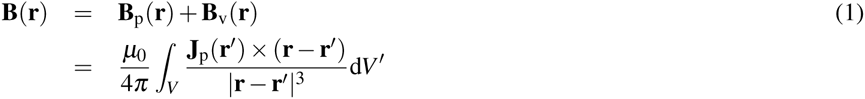

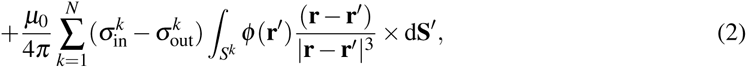

where **B**_p_(r) and **B**_v_(**r**) are the magnetic fields due to primary and volume currents, μ_0_ is the permeability of free space, and 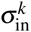, and 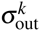 are the conductivities inside and outside the *k*th surface *S*^*k*^, respectively. The magnetic field can thus be obtained by first solving for the potential on conductivity boundaries and then integrating. Boundary element method (BEM) is a convenient choice for computing the potential and magnetic field in a piecewise homogeneous volume conductor (e.g. Stenroos et al. (2007)).

### 2.2 Forward model

The primary current distribution is typically discretized into a set of current dipoles (Hämäläinen et al., 1993). The output of a sensor is obtained by integrating the magnetic field due to primary-current dipoles through the sensitive volume of the sensor; the sensitive volume is discretized into a set of integration points and the integral is approximated by a weighted sum of the magnetic field components at these points.

With these models, the magnetic-field amplitudes at the *N*_*c*_ sensors 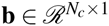 are related to the *N*_*s*_ amplitudes of the current dipoles 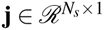 by a linear mapping

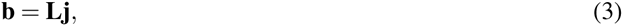

where 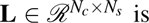 is the lead-field matrix. The ith column of the lead-field matrix represents the magnetic field pattern of the *i*th unit source, i.e., the topography of source (**t**_*i*_); the *j*th row represents the sensitivity pattern of the *j*th sensor to the sources, i.e., the lead field of sensor (**l**_j_). According to the Geselowitz formula presented earlier, the lead-field matrix can be considered a sum of two matrices:

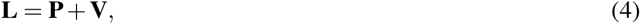

where **P** and **V** represent the contributions of the primary and volume currents, respectively.

### 2.3 Minimum-norm estimate

Minimum-*l*_2_-norm estimator (Dale and Sereno, 1993; Hämäläinen and Ilmoniemi, 1994) can be used to estimate underlying sources from measured data. The source estimate, which satisfies the regularized minimum-*l*_2_-norm condition and takes into account the spatiotemporal properties of the noise, is

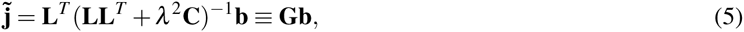

where λ^2^ is the regularization parameter, **C** is the noise covariance matrix and **G** is the minimum-norm estimator. The regularization parameter has been suggested to be chosen as (Lin et al., 2006a,b)

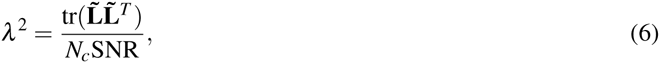

where *N*_*c*_ is the number of sensors, SNR is the estimated signal-to-noise ratio, tr(·) is trace of a matrix and L is the 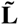 whitened lead-field matrix: 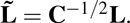.

## 3. Methods

Here, we present the methods and models that were used in the simulations and review the measures that we used to quantify the performance of the arrays. First, we constructed the anatomical models from the magnetic resonance images (MRIs), modeled the sensors and sensor arrays, and then computed the lead-field matrices. Prior to the actual simulations, we quantified whether the lead-field matrices of the OPM arrays were more sensitive to the skull-conductivity value and to the densities of BEM surface tessellations than the SQUID arrays. Subsequently, we built the lead-field matrices and minimum-norm estimators for the computation of performance metrics.

### 3.1 3.1. Anatomical models

T1-weighted MRIs were obtained from ten healthy adults (seven males, three females) using a 3-T scanner and a 3D MP-RAGE sequence. FreeSurfer software (Dale et al., 1999; Fischl et al., 1999a; Fischl, 2012) was used to segment the cortical mantles from individual MRIs. The primary current distributions were assumed to lie on the cortical surfaces and were discretized to sets of dipoles oriented normal to the local cortical surface (10 242 dipoles per hemisphere). For each subject, we used the watershed algorithm (Ségonne et al., 2004) implemented in FreeSurfer and MNE software (Gramfort et al., 2014) to segment the brain, skull and scalp compartments. We triangulated and decimated these surfaces to obtain three meshes (2 562 vertices per mesh) for the BEM. To estimate errors due to the coarser meshes, we constructed an additional model for one subject with surfaces comprising 10 242 vertices.

To obtain group-level metrics, the spherical morphing procedure in Freesurfer (Fischl et al., 1999b) was used to map the values from individual subjects to their average brain, where the values were subsequently averaged.

### 3.2 Sensor models

The sensor output was computed by integrating over a set of points within the sensing volume (OPMs) or pick-up coil plane (SQUIDs). The descriptions of the sensor geometries are presented in Table 1. The OPM sensor was a cube with a sidelength of 5 mm, and the integration points were distributed uniformly within that volume. The SQUID sensors were modeled as in the MNE software (Table 1; Gramfort et al. 2014). The OPMs were assumed to have a noise level of 6 fT/ 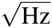 while the noise levels of SQUID magnetometers and gradiometers were 3 fT/ 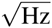 and 3 fT/ (cm 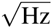), respectively.

**Table 1:** Sensor models used in this study. *N* is the number of integration points ***r***, *w* is the weight of each point and *σ* is the noise level.

| Sensor type | $N$ | $\mathbf{r}(x, y, z)$ [mm] | $w$ | $\sigma$ |
| --- | --- | --- | --- | --- |
| OPM | 8 | $(\pm 1.25, \pm 1.25, 1.25)$ | 1/8 | $6 \text{ fT}/\sqrt{\text{Hz}}$ |
| | | $(\pm 1.25, \pm 1.25, 3.75)$ | 1/8 | |
| SQUID gradiometer | 2 | $(\pm 8.4, 0, 0.3)$ | $\pm 1/16.8 \text{ mm}$ | $3 \text{ fT}/(\text{cm} \sqrt{\text{Hz}})$ |
| SQUID magnetometer | 4 | $(\pm 6.45, \pm 6.45, 0.3)$ | 1/4 | $3 \text{ fT}/\sqrt{\text{Hz}}$ |

### 3.3 Sensor arrays

The SQUID sensor arrays of the Elekta Neuromag^®^ MEG system (102 magnetometers; 204 planar gradiometers; Elekta Oy, Helsinki, Finland) were positioned around the subjects’ heads as in a real MEG measurement. For each subject, the position was determined so that the SQUID array covered the cortex uniformly and symmetrically along the left-right axis. Thereafter, the sensor positions were verified to be at least 2 cm (approximate thickness of the MEG dewar) from the scalp. The SQUID array for each subject is shown in Fig. 1. We considered three variants of this array: planar gradiometers only (gSQUID), magnetometers only (mSQUID), and the full 306-channel array (aSQUID) with both sensor types. Each of these arrays measures the amplitude and/or the gradient of the magnetic field component normal to the surface of the MEG helmet.

**Figure 1:**
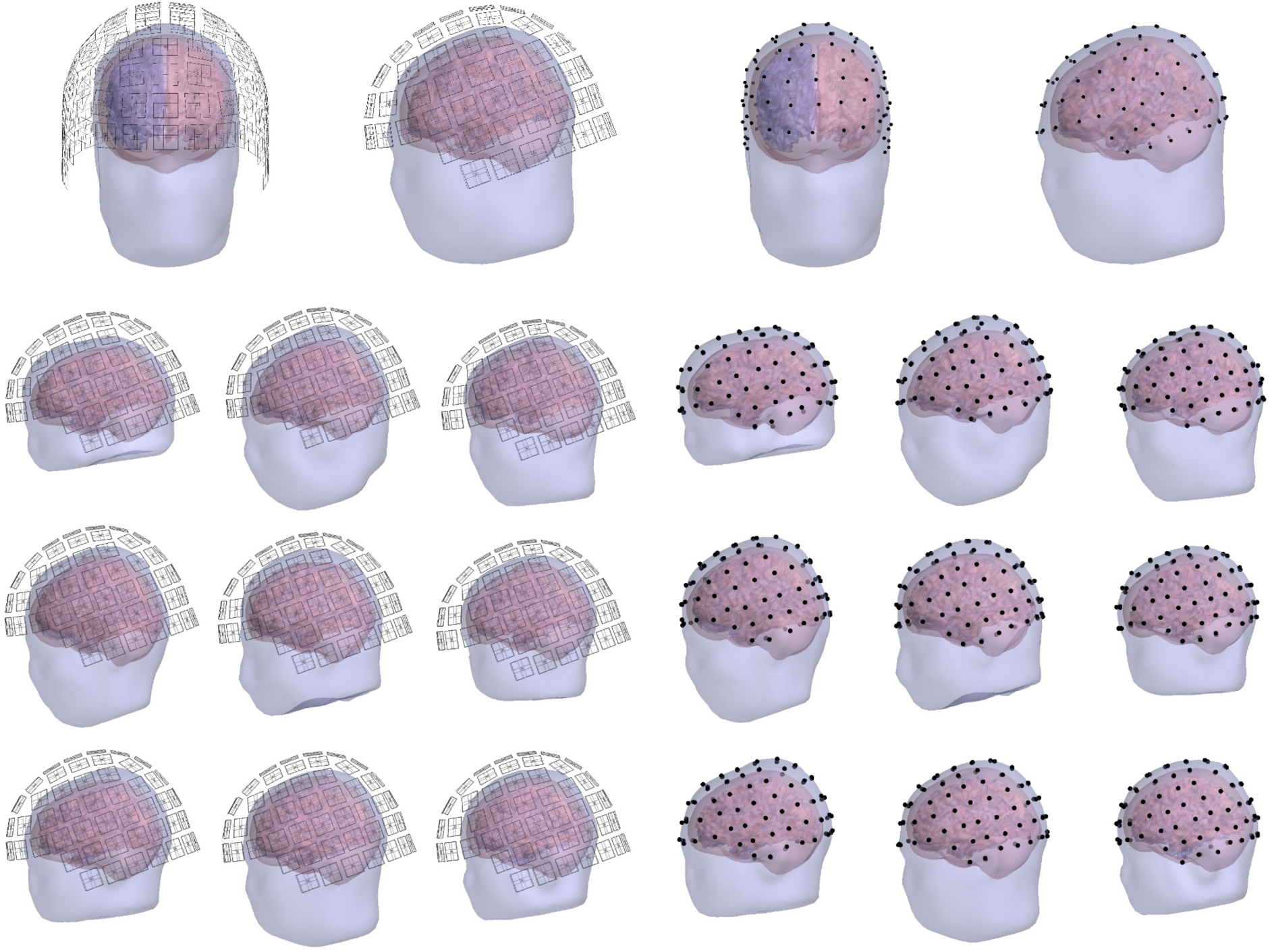
The constructed SQUID (left) and OPM (right) sensor positions. Top: Frontal and lateral views of the sensor locations for one subject. Other rows: Lateral views for the rest of the subjects.

The OPM locations (shown in Fig. 1) were derived from the SQUID arrays by projecting the sensor locations to the scalp. The distance of the closest face of the OPM sensor cube to scalp was set to 1 mm. For these positions, we defined three different OPM arrays: one with 102 sensors measuring the normal component of the magnetic field with respect to the local surface of the scalp (nOPM), other with 204 sensors measuring the two orthogonal tangential components (tOPM) and a combination of the aforementioned arrays, with 306 sensors measuring all field components (aOPM).

### 3.4 Forward models

For each subject, the lead-field matrix **L** was computed using a linear Galerkin BEM formulated with Isolated Source Approach (ISA) (Stenroos and Sarvas, 2012). The conductivities of the brain, skull and scalp compartments of the BEM model were set to [1 1/25 1]×0.33 S/m. To assess the sensitivity of the calculated lead-field matrices to skull conductivity, in one subject we compared the computed topographies against those obtained with skull conductivity values of 0.33/50 and 0.33/80 S/m.

### 3.5 Minimum-norm estimator

We estimated the SNR in the regularization parameter of Eq. (6) with the average SNR of the cortical sources; we defined the SNR of the *i*th source as the average SNR across the sensors

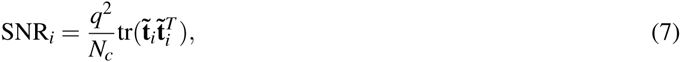

where *q*^2^ is the source variance and 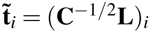 is the whitened topography of the *i*th source. We took into account only sensor noise which was assumed to be uncorrelated across the sensors; thus, we used a diagonal noise covariance matrix **C**, whose diagonal elements were the noise variances 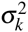 of the sensors. With a diagonal **C**, Eq. (7) simplifies into (Goldenholz et al., 2009)

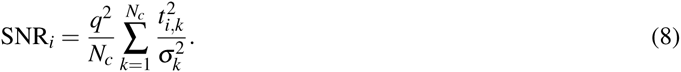

We set the source variance *q*^2^ so that the average SNR across the sources was 1 for the SQUID magnetometers and calculated the average SNRs and regularization parameters for the other arrays using that particular source variance.

### 3.6 Metrics

We computed topography power of the arrays and analyzed the contributions of the primary and volume currents to the signals. We then assessed the correlations between source topographies and examined the lead fields of the sensors by comparing the fields-of-view (FOVs) of the sensors and calculating correlations among the lead fields. To complete forward model -based metrics and quantify the performance of the array with a single number, we calculated the total information conveyed by the array (Kemppainen and Ilmoniemi, 1989). Last, we investigated the point-spread functions (PSFs) of the arrays in the minimum-norm estimation. We calculated all metrics individually for each subject, morphed the results to the average brain and averaged.

*Signal power.* We defined the topography power of ith source as the *l*_2_-norm of the source topography squared, ∥**t**_*i*_∥^2^. If we assume that all sensors in the array have equal sensor noise variance σ^2^, topography power is linearly proportional to SNR of source as Eq. (7) simplifies to

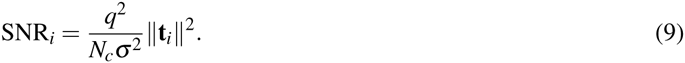

To compare the overall signal power of the arrays, we defined the relative power

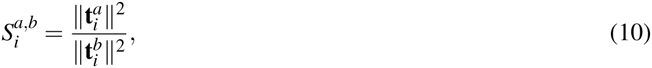

where 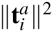 and 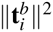 are the topography power of the source *i* in arrays *a* and *b*, respectively. Assuming again equal sensor noise variances in the arrays, the ratio of SNRs between arrays is

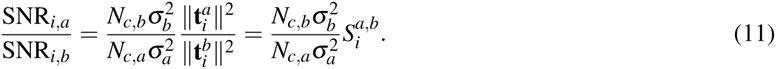

*Primary- and volume-current contributions*. We studied also the contributions of the primary and volume currents to the total magnetic field across the different sensor arrays. We defined TP as the ratio of the norms of the topographies of the total and primary current as

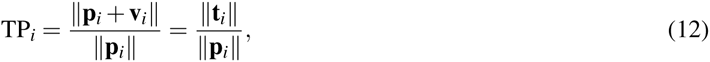

where **p**_*i*_, **v**_*i*_, and *t*_*i*_ denote the *i*th columns of the corresponding matrices **P, V** and **L**. Considering the values of TP, three different scenarios arise:

- TP < 1: Volume-current field decreases the overall amplitude of the primary-current field.
- TP **≈** 1: Volume currents do not noticeably decrease or increase the overall amplitude of the primary-current field.
- TP > 1: Volume-current field increases the overall amplitude of the primary-current field.

We also quantified the relative overall magnitude of the topographies of the primary and volume currents for the different sensor arrays by calculating the ratio of the norms of these field components

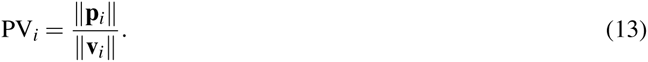

In addition to these amplitude-based measures, we investigated the differences in the field shapes of primary and volume currents by computing the correlation coefficient CC_PV_ between their topographies,

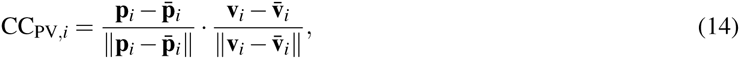

where · is the dot product and 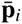 denotes the mean of **p**_*i*_.

*Topography overlap.* We calculated the correlation coefficient between the topography of the reference source and the topographies of all the other sources using

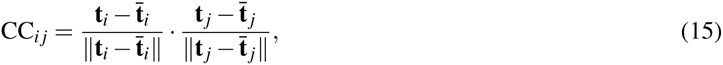

where *i* is the index of the reference source and *j* goes through the indices of all the other sources. By labeling those sources with high correlations (absolute value of CC ≥ 0.9), we estimated the peak position error (PPE) and cortical area (CA) of the reference source. PPE is the distance from the reference source to the center-of-mass of the highly-correlated sources; small values of PPE indicate that the correlated sources are scattered close to the reference source. CA is the relative cortical area of the highly-correlated sources and quantifies the spread of the sources that exhibit similar topographies; small values of CA indicate minor spread or a small number of correlated sources. We calculated CA as a percentage of the total area of the cortex that is spanned by the highly-correlated sources. These metrics have been previously used to assess localization accuracy (PPE) and spread (CA) of minimum-norm estimates (Stenroos and Hauk, 2013). PPE has also been used to assess the effect of the depth-weighting parameter to minimum-norm estimates (Lin et al., 2006b). PPE and CA are of the reference source *i* are expressed as

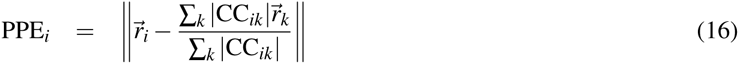

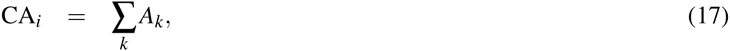

where the sums are taken across the labeled sources, CC_*ik*_ is the correlation coefficient between the topographies of the reference source *i* and source *k*, 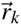 and 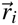 are the locations of source *k* and the reference source, respectively, and A_*k*_ is the relative cortical area associated with source *k*.

*Sensor lead fields.* We quantified the effect of sensor orientation and distance from the scalp on its FOV; for each sensor, we calculated the relative cortical area (Eq. (17)) for which the lead-field amplitude of the sensor was more than half of the maximum amplitude. This metric quantifies the effective field-of-view (eFOV) of the sensor as it measures the area spanned by the sources that are most visible to the given sensor. In addition, we investigated the overlap of the sensor lead fields as a function of the sensor distance (de Munck et al., 1992): for each sensor in the array, we calculated the distances and lead-field correlations between that sensor and all other sensors in the array.

*Total information.* Previously, the performance of various MEG arrays has been assessed by computing the total information *I*_tot_ conveyed by the array (Kemppainen and Ilmoniemi, 1989; Nenonen et al., 2004; Schneiderman, 2014). *I*_tot_ quantifies all of the aspects of the forward-model-based metrics, e.g., sensor distances to the sources, sensor configuration, sensor type, SNR and dependencies of the sensor lead fields, with a single number. We also used this metric to evaluate the performance of the arrays.

We assume that the signal and the noise of a channel are independent and normally distributed; according to Shannon’s theory of communication the information per sample of ith channel is then 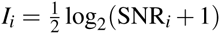, where SNR_*i*_ is the power signal-to-noise ratio of the channel (Shannon and Weaver, 1949). We further assume that the source time-series are uncorrelated and that their amplitudes follow Gaussian distribution: *j*_*i*_ ~ *𝒩* (0,*q*^2^), where *q*^2^ assumes the same value as in Sec. 3.5; in addition, we only take into account the sensor noise. The SNR of channel *i* is 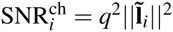, where 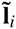 is the *i*th row of the whitened lead-field matrix 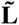. With the assumption of a diagonal noise covariance matrix, SNR simplifies into 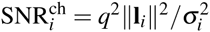.

To obtain the total information conveyed by the array by summing the contributions of individual channels, the sensor lead fields must be orthogonalized via an eigenvalue decomposition. First, we generate the matrix

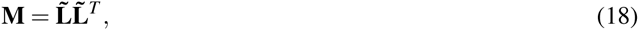

and compute its eigenvalue decomposition **M** = **USU***^T^*, where the columns of **U** are the eigenvectors of **M** and **S** is a diagonal matrix with eigenvalues λ_*j*_ of **M** in the diagonal. The orthogonalized lead fields are then 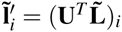 while the orthogonalized SNRs of the channels are 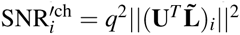. The total information per sample of a multichannel system is then given by

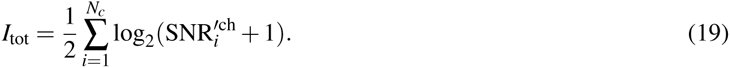

*Point-spread functions.* To evaluate the differences between the arrays in linear distributed-source estimation, we used metrics derived from the resolution matrix (Hauk et al., 2011; Liu et al., 2002; Molins et al., 2008; de Peralta Menendez et al., 1997; Stenroos and Hauk, 2013). Resolution matrix **K** gives the relation between the estimated and modeled current distributions as follows:

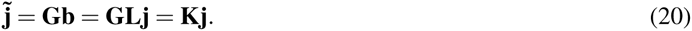

The columns of **K** are PSFs and they describe how a point source is distorted by the imaging system. By replacing CC in Eqs. (16) and (17) by PSF and picking the indices for which |PSF_*i*_| ≥ 0.5×PSF_*max*_, we can quantify the accuracy of a sensor array in minimum-norm estimation with PPE and CA; PPE quantifies the localization accuracy while CA estimates the spread of the PSF.

## 4. Results

### 4.1 Forward metrics

The relative powers for different combinations of nOPM, tOPM and mSQUID arrays are illustrated in Fig. 2. On average, the topography powers of the nOPM and tOPM are 7.5 and 5.3 times higher than the topography power of mSQUID. The topography powers of the OPM arrays vs. mSQUID are still higher for the superficial sources (~9.4 for nOPM and ~7.1 for tOPM) and decrease for deeper sources (~6.5 for nOPM and ~4.5 for tOPM). The topography power of the nOPM array is on average 1.5 times higher than that of tOPM, and it is higher for almost every point of the source space.

**Figure 2:**
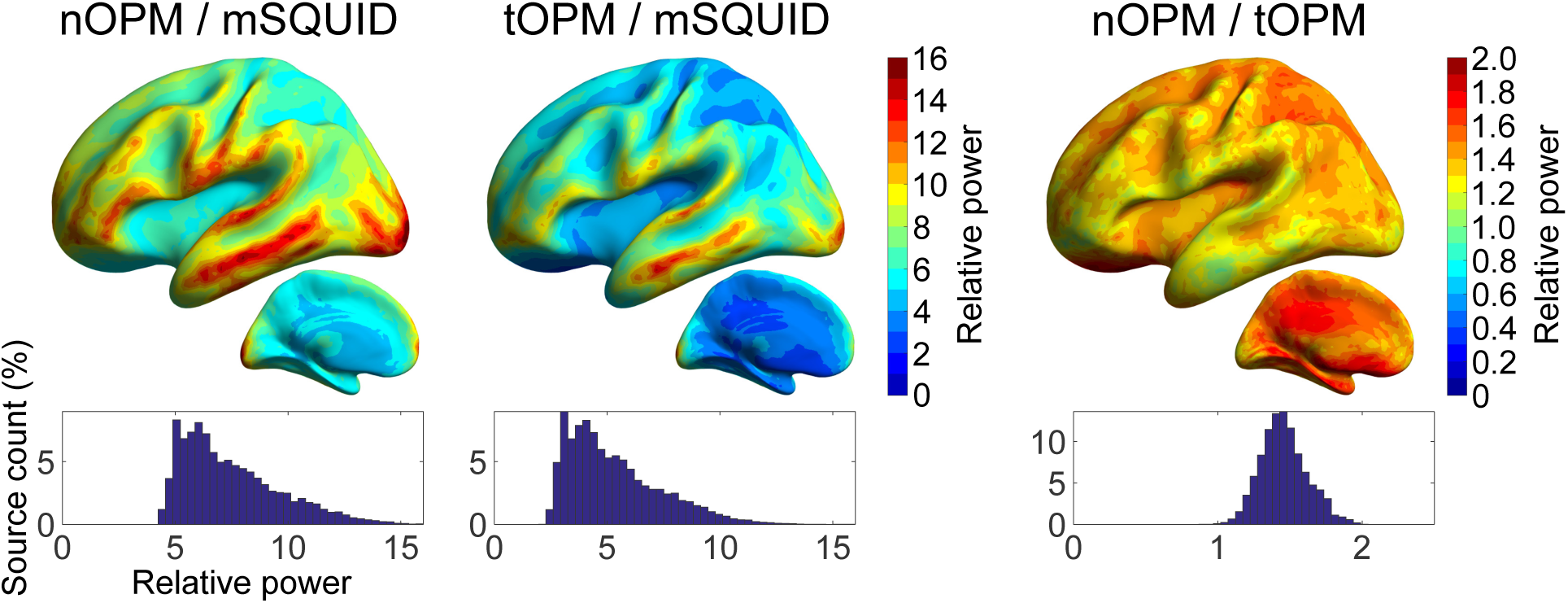
The relative powers of the arrays. The average values of the relative power across ten subjects are presented in the average brain of the subjects. Histograms represent the distribution of the values of relative power for sources in both hemispheres. Histograms are normalized by the total number of sources.

The comparisons of the primary- and volume-current components of the topographies are displayed in Fig. 3. The overall amplitudes of the volume current topographies are of the same magnitude as those of the primary currents for tOPM as the values of PV are concentrated around one. The field due to volume currents substantially suppresses the overall amplitude of the primary-current topography in tangential measurements since TP << 1 for tOPM (mean 0.28, range 0.05-0.46). For nOPM and mSQUID, the overall magnitude of the primary current topography is much higher than that of volume currents, as the majority of PV >> 1. Additionally, volume currents do not result in a major decrease in the overall amplitude of primary current topography in nOPM and mSQUID arrays as the values of TP are closer to one (mean 0.90 and 0.85 for nOPM and mSQUID, respectively). Volume currents increase the visibility of some deep sources in particular when measuring with nOPM and mSQUID arrays (regions where TP > 1). Furthermore, the closer the normal-component measuring sensors are to the sources, the larger the overall magnitude of the primary-current topography is relative to the volume-current topography since the mean and range of PV are 3.41 and 0.71–6.41 for nOPM and 2.67 and 0.62–4.81 for mSQUID. As the majority of the CC_PV_ values are well below zero, the volume-current topographies tend to have the opposite shape to the primary-current topographies. For tOPM, the correlation coefficients are very close to -1, i.e., exactly opposite topographical shape of the volume currents compared to the primary currents. The average CC_PV_ for nOPM, tOPM and mSQUID is -0.44, -0.95 and -0.53 while the ranges are -0.94-0.23, -1.0-(-0.89) and -0.97–0.17, respectively.

**Figure 3:**
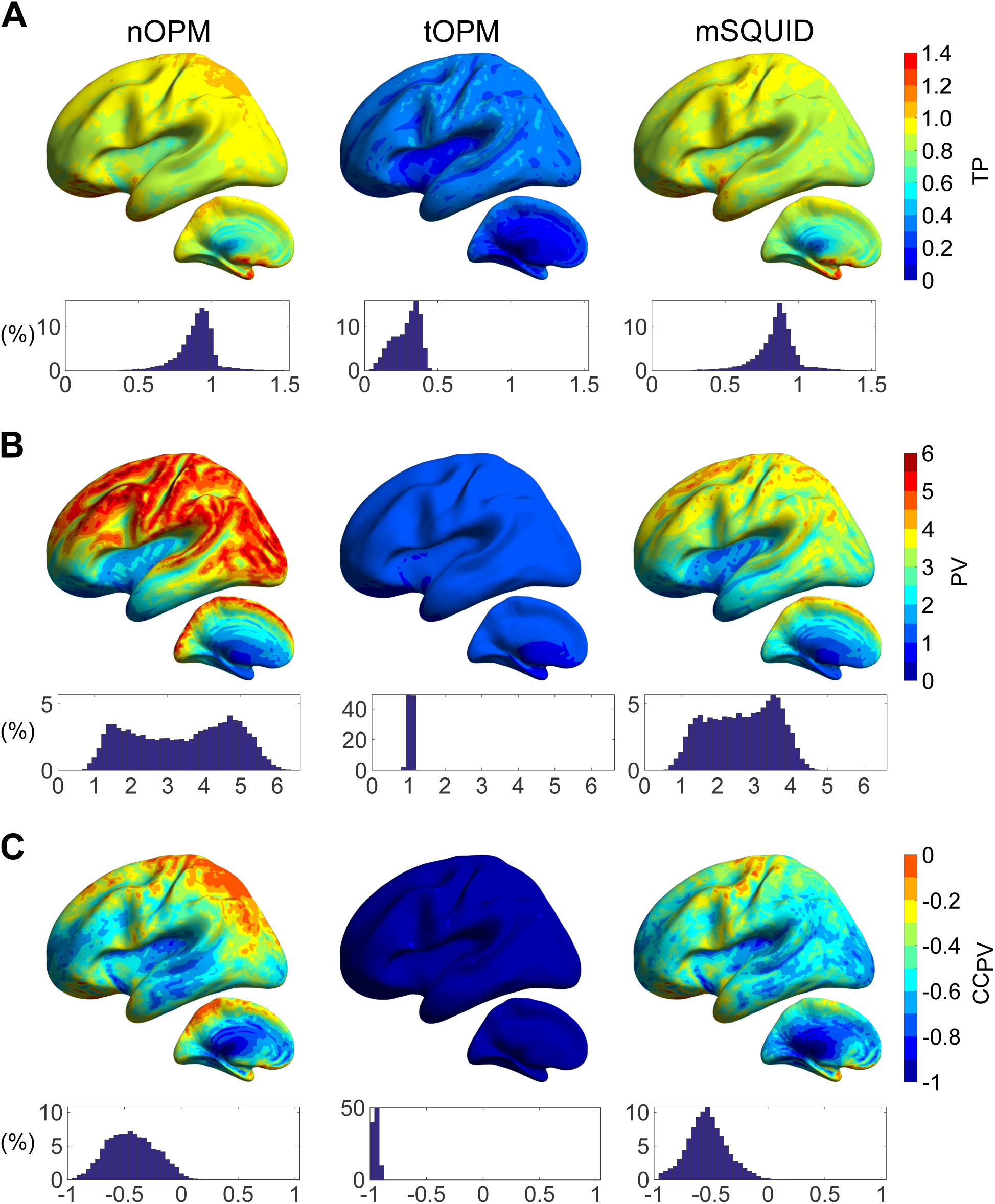
Contributions of primary and volume currents. The ratios of the norms of (A) total- and primary-current topographies (TP) and (B) primary- and volume-current topographies (PV). C: The correlation coefficient between the topographies of the primary and volume currents (CC_PV_). For further details, see the caption of Fig. 2.

Fig. 4 shows the peak position error (PPE) and cortical area (CA), which were based on correlations between the source topographies. CA is clearly smaller for nOPM, tOPM and gSQUID than for mSQUID, indicating that the topographies of the sources show less overlap in OPM and SQUID-gradiometer arrays. PPE is also smaller for the nOPM, tOPM and gSQUID suggesting that the similar-topography sources are closer to the actual source for these arrays. The measures are similar between nOPM, tOPM and gSQUID. The averages for PPE are 0.51, 0.45, 0.79 and 0.52 cm and for CA they are 0.26%, 0.19%, 0.66% and 0.26% for nOPM, tOPM, mSQUID and gSQUID, respectively.

**Figure 4:**
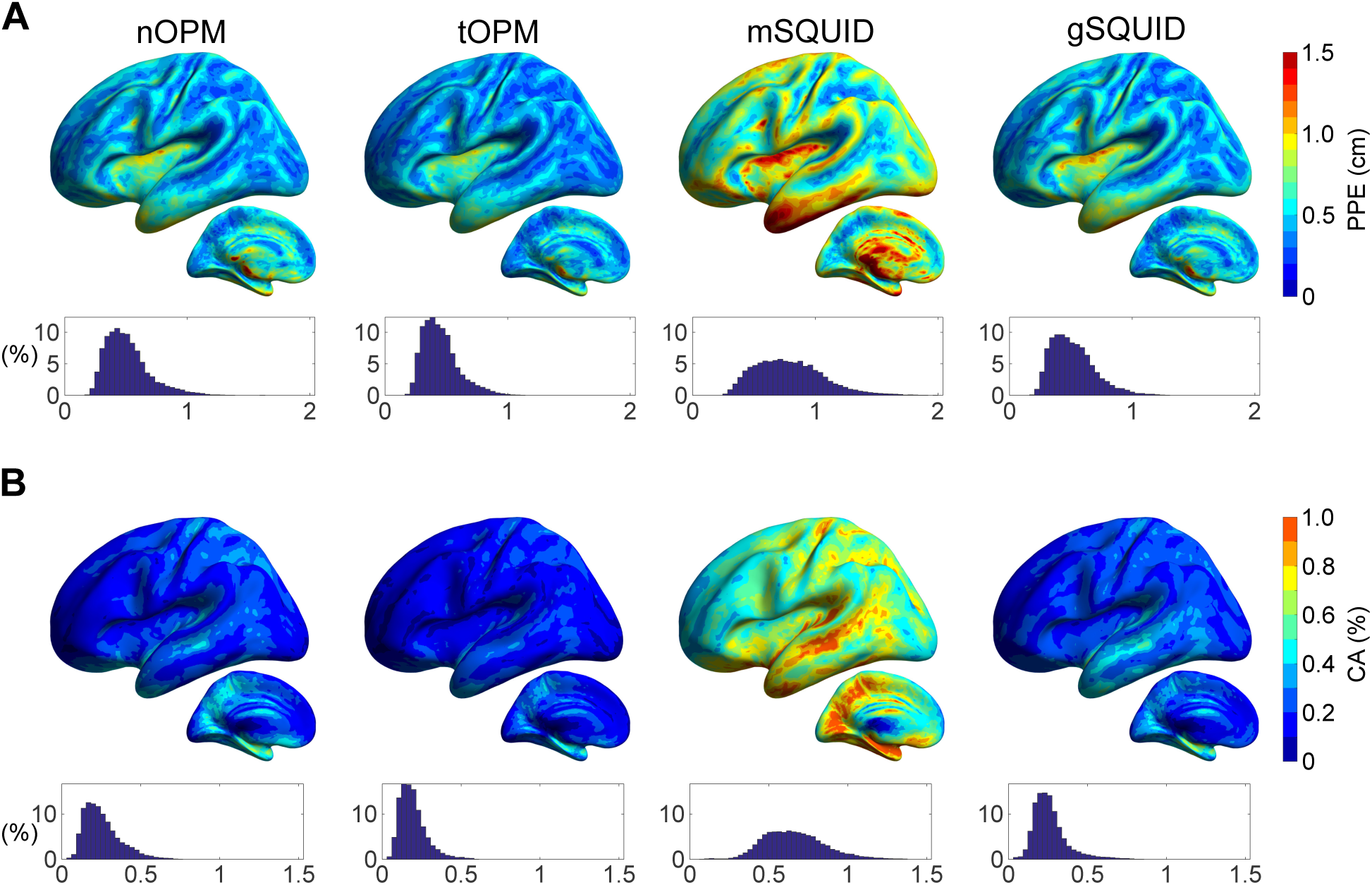
Similarity of source topographies. (A) Peak position errors (PPEs) and (B) cortical areas (CAs) of topography correlations. CA is measured as a percentage of the total area of cortex. For details, see the caption of Fig. 2.

The computed eFOVs of the sensors are illustrated in Fig. 5 as histograms into which the values from all subjects were pooled. When the magnetometers are brought closer to the scalp, their FOV shrinks as the values of eFOV are, on average, smaller for nOPM and tOPM than for mSQUID. Additionally, the eFOVs in tOPM are more focal than those in nOPM. The SQUID gradiometers also show smaller FOVs than the SQUID magnetometers. The eFOVs of gSQUID are similar to those of nOPM and tOPM. The mean values of eFOVs are 0.69%, 0.25%, 3.02% and 0.53% for nOPM, tOPM, mSQUID and gSQUID, respectively.

**Figure 5:**
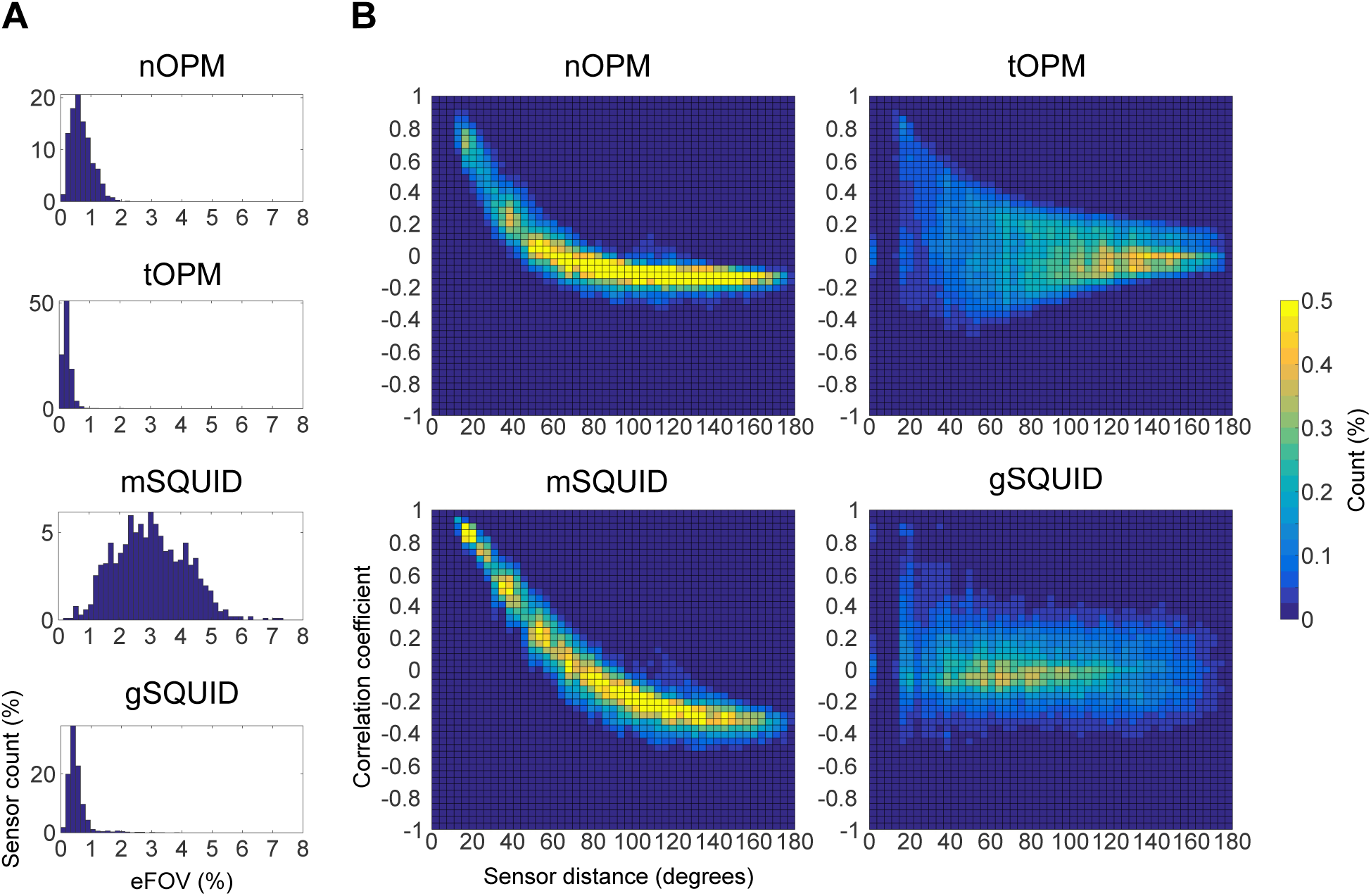
The effective field-of-view (eFOV) of the sensors and correlation of lead fields. A: Histograms of eFOV with values from different subjects pooled. eFOV is measured as a percentage of the total area of cortex. Each histogram has been normalized by the total number of sensors across the subjects. B: Histograms presenting the correlation coefficient between sensor lead fields as a function of sensor distance in degrees. The lead-field correlations across the subjects and sensors have been pooled. Each histogram has been normalized so that the sum over the bins gives 100%.

The lead-field correlations as a function of the sensor distance are presented in Fig. 5 as histograms into which the values from all subjects were pooled. The lead-field correlations in nOPM and mSQUID show a clear decreasing trend while the histograms of tOPM and gSQUID are more diffuse. Compared to mSQUID, the correlation between the lead fields of nOPM falls off faster as a function of distance. The correlations in tOPM exhibit a peak around zero distance since the lead fields of the co-located sensors that measure two orthogonal tangential components are nearly independent; the same applies also for gSQUID. The histograms also show that the number of strongly lead-field-correlated sensors is higher in mSQUID than in nOPM, tOPM or gSQUID.

The total information capacities of the arrays are presented in Fig. 6. Across the subjects, mSQUID and gSQUID convey similar amount of information. Combining mSQUID and gSQUID to aSQUID increases information capacity slightly. Both of the OPM arrays (nOPM and tOPM) provide more information than aSQUID. tOPM yields more information than nOPM and the combined array aOPM even more. The total information capacity per sensor is highest in nOPM.

**Figure 6:**
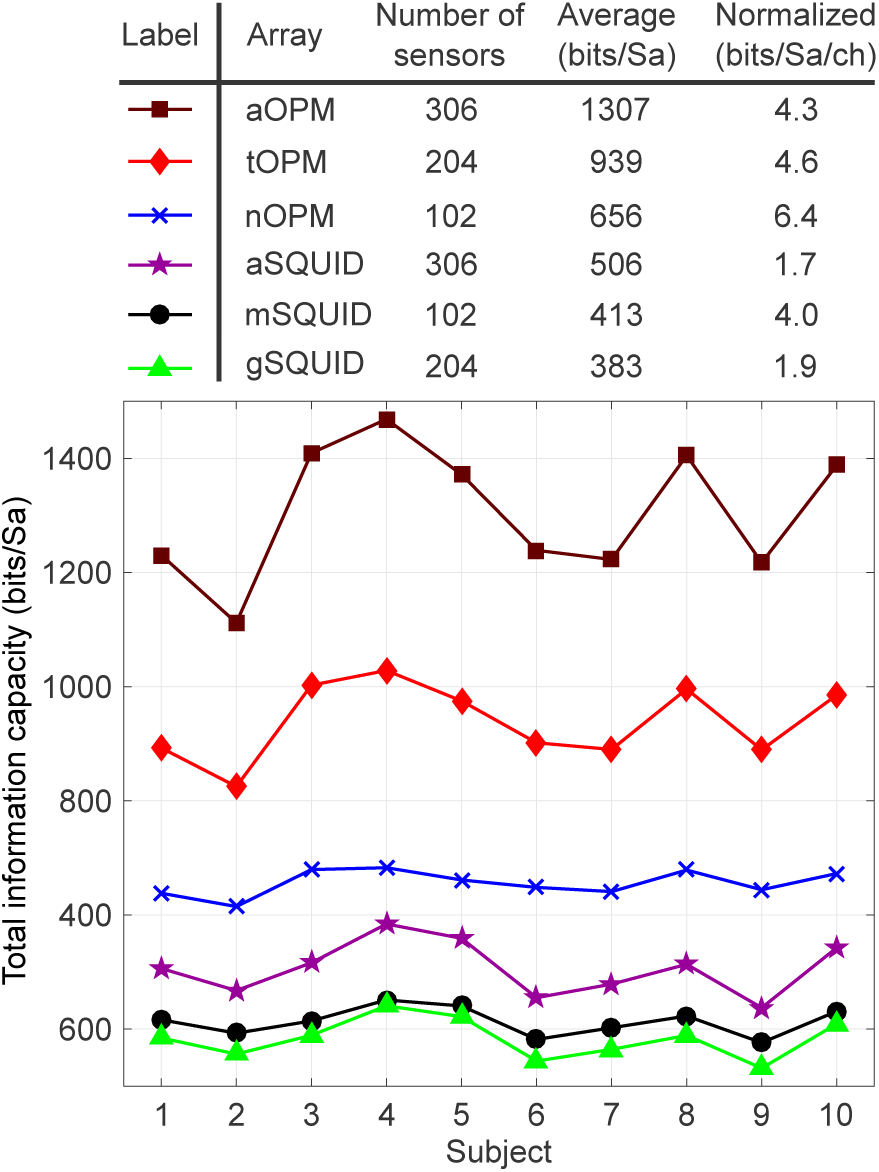
The total information capacities of the sensor arrays. Total information capacity is measured in units of bits per sample (bits/Sa). Top: Total information capacity averaged across the subjects and normalized by the number of sensors. Bottom: Total information capacity for each subject.

### 4.2 Point-spread functions

The results for the PSF-based metrics are displayed in Fig. 7 for nOPM, tOPM, aOPM, mSQUID and aSQUID arrays. Cortical maps and histograms of PPE between the different arrays exhibit similarities; yet, differences are evident in CA. Comparison between the normal-component-measuring arrays indicates that PSFs are less spread in nOPM: CA is larger for every source for mSQUID; the averages of CA are 0.67% and 1.57% for nOPM and mSQUID, respectively. The spread of the PSF does not necessarily correlate with the localization accuracy as the PPE of nOPM is larger for 60% of the sources. The averages of PPE are 1.30 and 1.27 cm for nOPM and mSQUID, respectively. For the most superficial sources the PPE is smaller in nOPM: averages of the PPE for the most superficial sources of left hemisphere are 0.48 cm for nOPM and 0.56 cm for mSQUID.

**Figure 7:**
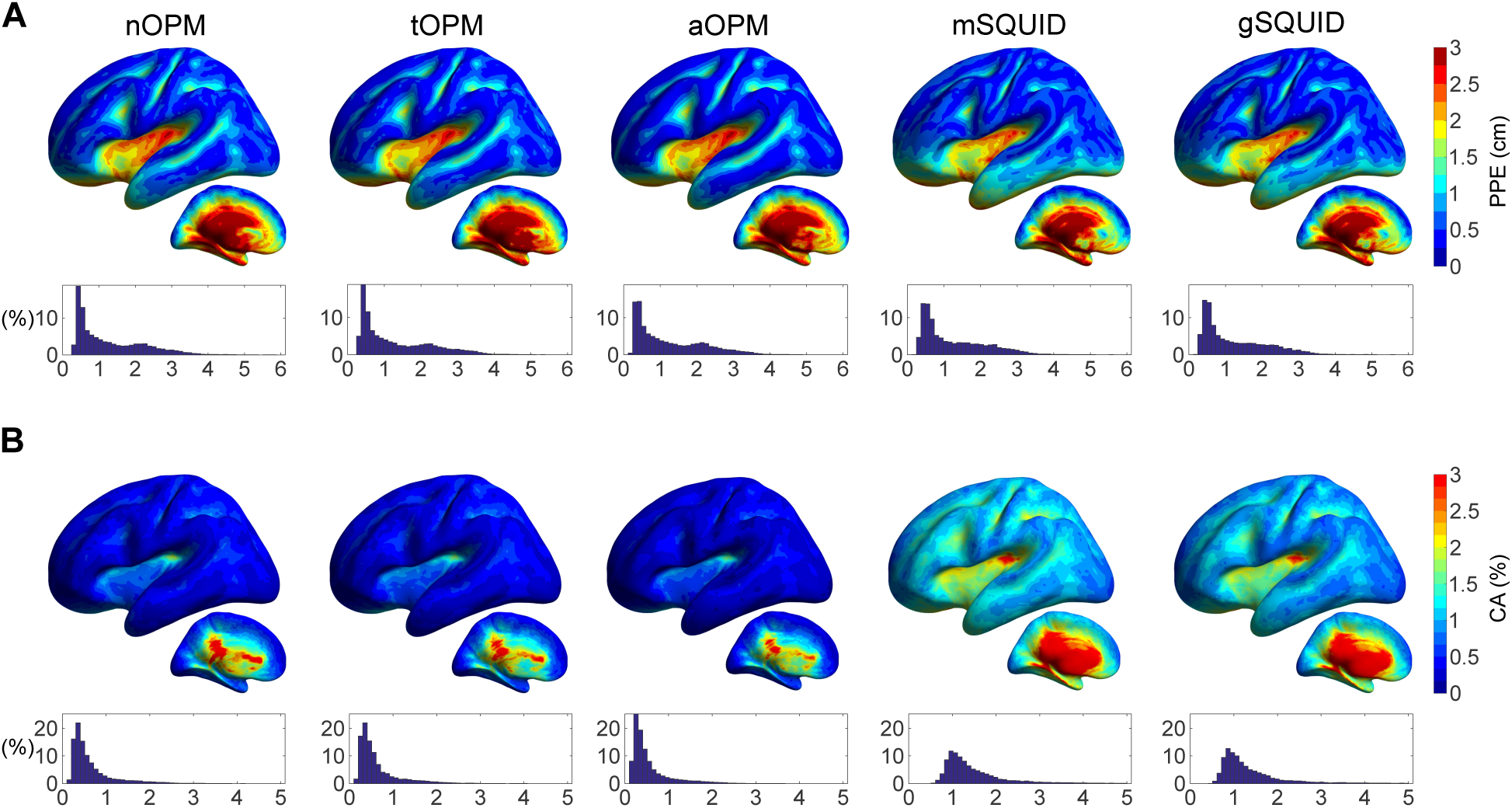
Characteristics of the minimum-norm-based point-spread functions. (A) Peak position error (PPE) and (B) cortical area (CA) relative to the total area of cortex. For details about the illustration, see the caption of Fig. 2.

The results between nOPM and tOPM are similar; averages of PPE and CA for tOPM are 1.31 cm and 0.64% while average PPE is 0.46 cm for superficial sources. For aOPM, the localization accuracy is slightly enhanced compared to the individual arrays, especially for the superficial sources. Averages of PPE and CA for aOPM are 1.26 cm and 0.57% for all sources while for the superficial sources the average PPE is 0.40 cm.

Comparison of the OPM arrays to the 306-channel SQUID system shows that there is a substantial decrease of the PSF spread for all OPM arrays while the localization performance (PPE) is similar. Nonetheless, for superficial sources the localization accuracy of aOPM is better than that of aSQUID; for superficial sources the average PPE of aSQUID is 0.53 cm. The averages of PPE and CA of aSQUID are 1.26 cm and 1.45%.

## 5. Discussion

The aim of this study was to assess the possible benefits of on-scalp MEG arrays that can be constructed using novel magnetic field sensors, e.g., optically-pumped magnetometers (OPMs) or high-T_c_ SQUIDs. The assessment was done by comparing the performance of state-of-the-art low-*T_c_* SQUID sensor arrays to hypothetical on-scalp arrays. We modeled three variants of SQUID arrays that measured the amplitude and/or the gradient of the magnetic field component normal to the surface of the MEG helmet (mSQUID: 102 magnetometers; gSQUID: 204 planar gradiometers; aSQUID: combination of mSQUID and gSQUID). As hypothetical on-scalp arrays, we defined three OPM arrays that measured normal (nOPM; 102 sensors), tangential (tOPM; 204 sensors) or normal and tangential (aOPM; 306 sensors) components of the magnetic field with respect to the local surface of the scalp. In the comparison, we used metrics derived from both forward and inverse models that quantified signal power, information, similarity of source topographies, overlap of the sensor lead fields, localization accuracy and resolution. Although we refer to the on-scalp sensors as OPMs, the results are applicable to any kind of sensor with similar characteristics.

Recently, Boto and colleagues (2016) reported a simulation study where they investigated the performance of on-scalp MEG sensor arrays. In the simulations, they employed a SQUID sensor configuration of 275-channel axial gradiometer system, that measured the gradient of the field component normal to the MEG helmet, and determined the OPM sensor positions by projecting the SQUID locations to 4 mm from scalp. Their OPM array measured the magnetic field component normal to the scalp. In the following, we compare our results to theirs whenever possible.

### 5.1 Amplitude measures

Among the nOPM, tOPM, and mSQUID arrays, nOPM had the strongest overall topography power despite that the number of sensors in tOPM was double and the topography powers were not normalized by the sensor count; the average relative powers were 7.5 and 5.3 for nOPM/mSQUID and tOPM/mSQUID, respectively. Assuming noise levels of 6 and 3 fT/ 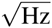 for OPM and SQUID magnetometers, respectively, the relative powers transform into overall signal-to-noise-ratio (SNR) gain of 1.9 for nOPM compared to mSQUID, while this figure is 0.7 for tOPM. For sources in the superficial parts of the brain, the average SNR of nOPM and tOPM are 2.4 and 0.9 times that of mSQUID. On the other hand, for equal overall SNR between the OPM and mSQUID arrays, the sensor noise levels should be 8.2 and 4.9 fT/ 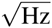 for nOPM and tOPM, respectively.

Boto and colleagues (2016) reported a fivefold SNR (amplitude) improvement with an OPM compared to a SQUID array, which was obtained by assuming equivalent sensor noise levels for the two arrays which both comprised axial gradiometers with 5-cm baselines. The authors computed the magnetic field with local-spheres head model (Huang et al., 1999) in a single head geometry. The position of the SQUID array with respect to the head was taken from a real MEG measurement where the back of the subject’s head was touching the helmet leaving a larger gap in the front. Thus, the values of relative SNRs were smaller (below four) for the occipital and larger for frontal sources (up to ten). We obtained average power-SNR gain of 7.5 (relative power between nOPM and mSQUID); thus, the average amplitude-SNR is approximately 2.7. The difference between the results can be largely explained by the different OPM and SQUID configurations (axial gradiometers vs. magnetometers) and helmet positionings. We located the SQUID arrays so that they offered uniform coverage of the cortex and symmetry along the left-right axis, thus yielding more uniform spatial distributions of relative power (Fig. 2). It is also possible that the head used in the simulations by Boto and colleagues was smaller than our average head size. The different volume conductor modelling approaches may also play a role (local spheres vs. three-shell BEM).

Our results suggest that in tOPM measurements, the field due to volume currents has a large screening effect on the field of the primary current. The shapes of the primary- and volume-current topographies in tOPM measurements are almost opposite for each source position as the average correlation coefficient (CC_PV_) between them is -0.95. In addition, as the average amplitude ratio between the primary- and volume-current toporaphies (PV) is 1.06, these topographies have very similar overall magnitude. Complete cancellation of primary-current topographies does not occur as CC_PV_ and PV are not exactly –1 (range –1.0–(–0.89)) and 1 (range 0.85–1.21), respectively. The ratio of overall amplitudes of total- and primary-current topographies (TP) is then larger than zero (average 0.28 and range 0.05–0.46).

The volume-current fields are also present in the normal component; they reduce the signal amplitudes of primary currents on average by 10% (nOPM array; calculated as (1 - TP) × 100%). This effect is much more drastic in measurements of the tangential component (tOPM array) where the overall signal reduction is 72%. Thus, measuring and modeling tangential field components is likely more sensitive to errors and simplifications in volume conductor models. In addition, since the relative primary-current contribution further increases when the normal-component-measuring sensors are brought closer to the scalp as the averages of PV are 3.4 for nOPM and 2.7 for mSQUID while the overall signal reductions are 10% and 15%, these sensors appear the optimal choice for on-scalp measurements.

### 5.2 Correlation metrics

We quantified the overlap of the source topographies using cortical area (CA) and peak position error (PPE): CA quantifies the area spanned by the similar-topography sources and PPE quantifies the relative locations of these sources. We considered source topographies similar if their correlation was over 0.9. To assess the sensitivity of CA and PPE to the choice of this threshold, we calculated CA and PPE using also thresholds 0.8, 0.85 and 0.95. The choice of the threshold did not substantially affect the relative measures between the sensor arrays.

CA and PPE showed that the OPM arrays offer clear benefits over SQUID magnetometers when it comes to the overlap of the topographies: the OPM arrays exhibit less-correlated topographies, and the correlated sources are closer to the actual sources which generate the topographies (see Fig. 4). In addition, these measures were slightly smaller (better) for the tOPM than for the nOPM, which can be due to the larger number of sensors in tOPM; more sensors allow more possibilities for the topographies to deviate from each other. The measures were smaller for gSQUID compared to mSQUID, showing the benefit of the focal sensitivity of planar gradiometers. In addition, the measures were similar between gSQUID and OPM arrays. Future studies could investigate whether the topography correlations can be further reduced by gradiometrization of OPMs.

The on-scalp OPMs showed more focal fields-of-view (FOVs) than the traditional SQUID magnetometers as the average values of the effective field-of-view (eFOV), which we defined as the percentage of the area of the cortex that is most visible to the given sensor, were 0.7%, 0.3% and 3.0% for nOPM, tOPM and mSQUID, respectively. Furthermore, the tangential vs. normal OPMs have more focal lead fields. The planar gradiometers also have smaller FOVs than the magnetometers at the same distance from the scalp; values of eFOV for gSQUID (average 0.53%) are of the same order as for the OPMs.

Lead-field correlations measure the degree of independency of the signals of the sensors in the arrays: the less correlated the lead fields are, the more independent measurements the array yields, and the more information is conveyed by the array. Sensors in nOPM, tOPM and gSQUID provide mutually more independent measurements than sensors in mSQUID as the OPM arrays and gSQUID had less lead-field-correlated sensors than mSQUID (see Fig. 5). The lead-field overlap fell faster as a function of the sensor distance in nOPM than in mSQUID: for a similar level of lead-field correlations in nOPM as in mSQUID, the number of OPM sensors could be much higher than 102. The faster decrease of the lead-field correlations in nOPM can be explained by the smaller sensitive elements and eFOVs of the OPMs and higher spatial frequencies available in the on-scalp measurements. In gSQUID, the correlation histograms peaked at zero correlation at almost every sensor distance (Fig. 5) demonstrating the benefits of planar gradiometers: due to more focal FOVs, the lead fields have less overlap. In addition, the co-located planar gradiometers have orthogonal lead fields. The results also show that the lead fields of sensors that measure orthogonal tangential components of the magnetic field have nearly independent lead fields.

### 5.3 Total information and point spread

According to our results, the information capacities of the OPM arrays are clearly higher than that of the state-of-the-art SQUID array. In addition, the tOPM array with 204 sensors conveys more information than the nOPM array with 102 sensors, which can be attributed to the larger number of sensors in tOPM; we verified that the measurement of only one tangential component (longitudinal or latitudinal) at 102 locations does not yield more information than the measurement of the normal component. Also the total information capacity per sensor is highest in nOPM.

Furthermore, our results clearly indicate that the normal and tangential components carry independent information as the information capacity of the aOPM is higher than the information capacities of its sub-arrays nOPM and tOPM, indicating that these measurements are not redundant; see also de Munck and Daffertshofer (2012). Similar observations have been made for cardiomagnetic fields (Arturi et al., 2004). To support the total information analysis, we provide the singular values of the lead-field matrices in Appendix A.

The results from the PSF analysis showed that localization accuracies of the OPM and SQUID arrays in minimum-norm estimation are similar. However, the OPM arrays provided substantially more focal point-spread functions and thus should offer higher spatial resolution. Yet, inverse methods that fully utilize the advantages of on-scalp measurements could be developed.

### 5.4 Models and field computation

We verified our three-shell models for numerical accuracy and sensitivity to parameters by comparing topographies obtained with different model parameters. We first computed the magnetic fields with BEM surfaces consisting of 2 562 vertices per boundary (average sidelength 7 mm in the scalp mesh). As the coarseness of the meshes could affect the results especially when fields are evaluated close to the surfaces, we repeated in one subject the computations using surfaces consisting of 10 242 vertices. The results suggest that the sensitivities of topographies to the densities of the BEM surface tessellations are of the same order of magnitude for OPM and SQUID arrays, when LG BEM is used, and that meshes with 2 562 vertices with a linear Galerkin (LG) BEM are sufficient for the OPM arrays. For skull conductivity, we used a value of 0.33/25 S/m. To assess, whether OPMs are more sensitive to this model parameter than SQUIDs, we compared the obtained topographies to those obtained with skull conductivity values of 0.33/50 and 0.33/80 S/m in one subject and found that the topographies were robust to the choice of the conductivity value (results not shown). This simulation with three-shell conductor models suggests that the OPM arrays are not more sensitive than the SQUID arrays to conductivity errors in a layer-like skull.

We also compared the topographies obtained with a linear collocation (LC) (De Munck, 1992) and LG BEM (Mosher et al., 1999) across the subjects. The errors were similar for nOPM and mSQUID arrays whereas for tOPM they were somewhat larger (relative difference of 0.4% vs. 4%). Thus, the field computation for normal-component-measuring OPMs could potentially be done with a LC BEM, which is computationally lighter than the more accurate solvers such as LG BEM or symmetric BEM (Kybic et al., 2005). OPMs that measure tangential components may require more accurate BEM solvers than LC BEM or, optionally, finer meshing.

The higher sensitivity of tangential-component-measuring OPMs to volume currents implies that these sensors are likely more sensitive to errors and simplifications in the volume conductor model, e.g., surface segmentation errors and the amount of detail modeled. As the volume-current sensitivities of nOPM and mSQUID are similar, the results of forward-modeling studies for SQUID arrays are probably valid for nOPM arrays, too. The effect of common volume-conductor-model errors and simplifications on both nOPM and tOPM could be assessed by comparing geometrically correct and erroneous simplified models to more detailed reference models (Stenroos et al., 2014; Stenroos and Nummenmaa, 2016; Vorwerk et al., 2014).

The sensing volume of an OPM sensor was modeled as a (5-mm)^3^ cube with eight integration points while the SQUID models were based on those in the MNE software. We assumed a noise level of 6 fT/ 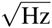 for the OPM sensor based on the work by Shah and Wakai (2013) who reported such a field resolution for their sensor in optimal conditions. Their sensor comprised a (4-mm)^3^ vapor cell, which is roughly equal to the sensitive volume in our OPM model. For the SQUID magnetometers and gradiometers, we assumed sensor noise levels of 3 fT/ 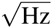 and 3 fT/cm 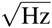, which are typical values for the Elekta Neuromag^®^ MEG systems.

We assumed the closest face of the OPM vapor cell to be 1 mm from the scalp; the distance between the scalp and the center of the sensitive volume is thus 3.5 mm. In practice, the thermal insulation needed between the outside surface of the sensor and the heated vapor cell imposes a lower limit on this distance. Alem and colleagues (2015) have manufactured OPMs with small vapor cells of (1.5 mm)^3^ and with a distance of 4.5 mm from the center of the cell to the outer surface of the sensor housing. OPMs by QuSpin Inc. (Louisville, USA) contain vapor cells of (3 mm)^3^ and exhibit a distance less than 6 mm from the center to the surface of the sensor. Thus, compared to the state-of-the-art OPMs, our hypothetical OPMs are optimistic in design; however, we anticipate that the distance between the vapor cell and the outer surface of the sensor will shrink in the next generation OPMs due to advances in sensor packaging and in the utilization of microfabrication (Liew et al., 2004).

### 5.5 Normal *vs.* tangential sensors

MEG devices have traditionally comprised normal-component-measuring sensors. Switching to a tangential array would introduce a profound change in the interpretation of the sensor-level data; e.g. the isocontours look very different in sensor arrays that measure either normal or tangential field components. In addition, compared to normal-component measurements, the higher sensitivity of tangential measurements to volume currents might necessitate more accurate head models and numerical methods (see Fig. 3 and discussion in Sec. 5.4). Individual differences in the conductivity profile of the head would render comparisons of sensor-level data between individuals less accurate in tangential-component-measuring sensor arrays as the data is more affected by the volume currents. Further, to obtain similar SNRs, the requirements for the sensor noise level are more strict for tangential sensors due to their lower topography power (see discussion in Sec. 5.1). For these reasons, we consider the normal-component-measuring sensors to be the optimal choice for MEG.

### 5.6 Implications to the construction of OPM arrays

Total information and topography-similarity measures offer means for designing optimal on-scalp sensor arrays. The benefit of optimizing an array with respect to these measures over, e.g., source-localization performance is that these measures depend only on the forward model and are thus general whereas the different source-localization methods may yield different optima. Higher information content should always benefit source localization; the task is then to develop localization methods that can fully exploit the additional information.

Since the scalp surface area is limited, adding more sensors to the array is beneficial only up to a point after which the utility of additional sensors decreases (Kemppainen and Ilmoniemi, 1989). After this point the lead fields of the sensors start to overlap so strongly that adding sensors no longer provide more information about the sources. The on-scalp arrays will benefit from a larger number of sensors than 102 and a denser array than mSQUID since the lead-field correlations decrease faster in nOPM than in mSQUID (Fig. 5).

Total information conveyed by the OPM arrays could likely be increased even further by adding OPM planar gradiometers in the arrays as the information conveyed by mSQUID can be increased with gSQUID (see Fig. 6). The baselines and exact types of the gradiometers could be deduced by simulating the total information capacity.

Sensor arrays that have minimal lead-field overlap and focal FOVs of the sensors offer also other advantages such as minimization of “field spread” (interdependencies of sensor signals), which would be particularly beneficial for functional connectivity analysis in the sensor space (see Schoffelen and Gross (2009)) and for the interpretability of sensor-level data.

Our total information analysis provides also a useful simulation benchmark. To facilitate the comparison of different sensor arrays, the source amplitudes can be set such that the average SNR across the sources for the SQUID magnetome-ters of the Elekta MEG system is one. Then, the SQUID magnetometers can be taken as a reference and metrics computed for other arrays using the same source amplitudes.

### 5.7 Future directions

We propose three directions for the development and establishment of on-scalp MEG. First, a thorough analysis of modeling errors on on-scalp arrays should be performed. In particular, it should be assessed whether the tangential-component-measuring OPMs provide added value due to their larger sensitivity to volume currents. Second, the on-scalp OPM array should be optimized for the number and types of sensors (magnetometers and gradiometers). This optimization can be performed with total-information analysis that is assisted with performance analysis of several source-localization methods. Also aspects on the interference rejection of the sensor array must be included in the design. Third, source-localization and interference-rejection methods that maximize the potential of on-scalp MEG arrays should be developed.

## 6. Conclusions

We examined the performance of hypothetical on-scalp MEG arrays compared to a current state-of-the-art MEG system. The results indicate that on-scalp arrays should offer clear benefits over traditional SQUID arrays in several aspects of performance; signal-to-noise ratio, total information conveyed by the array, and the achievable spatial resolution should improve substantially. These measures can be used to guide the design of on-scalp MEG arrays for optimal performance.

## 7. Acknowledgements

The authors thank Dr. Jukka Nenonen for comments on the manuscript, Advanced Magnetic Imaging centre of Aalto Neuroimaging Infrastructure for the MR-images, and Aalto University Science-IT for providing computational resources. Research reported in this publication was supported by the European Union FP7 project “Magnetrodes” (grant number 600730) and by the National Institute Of Neurological Disorders And Stroke of the National Institutes of Health under Award Number R01NS094604. The content is solely the responsibility of the authors and does not necessarily represent the official views of the funding organizations.

**Figure A.8:**
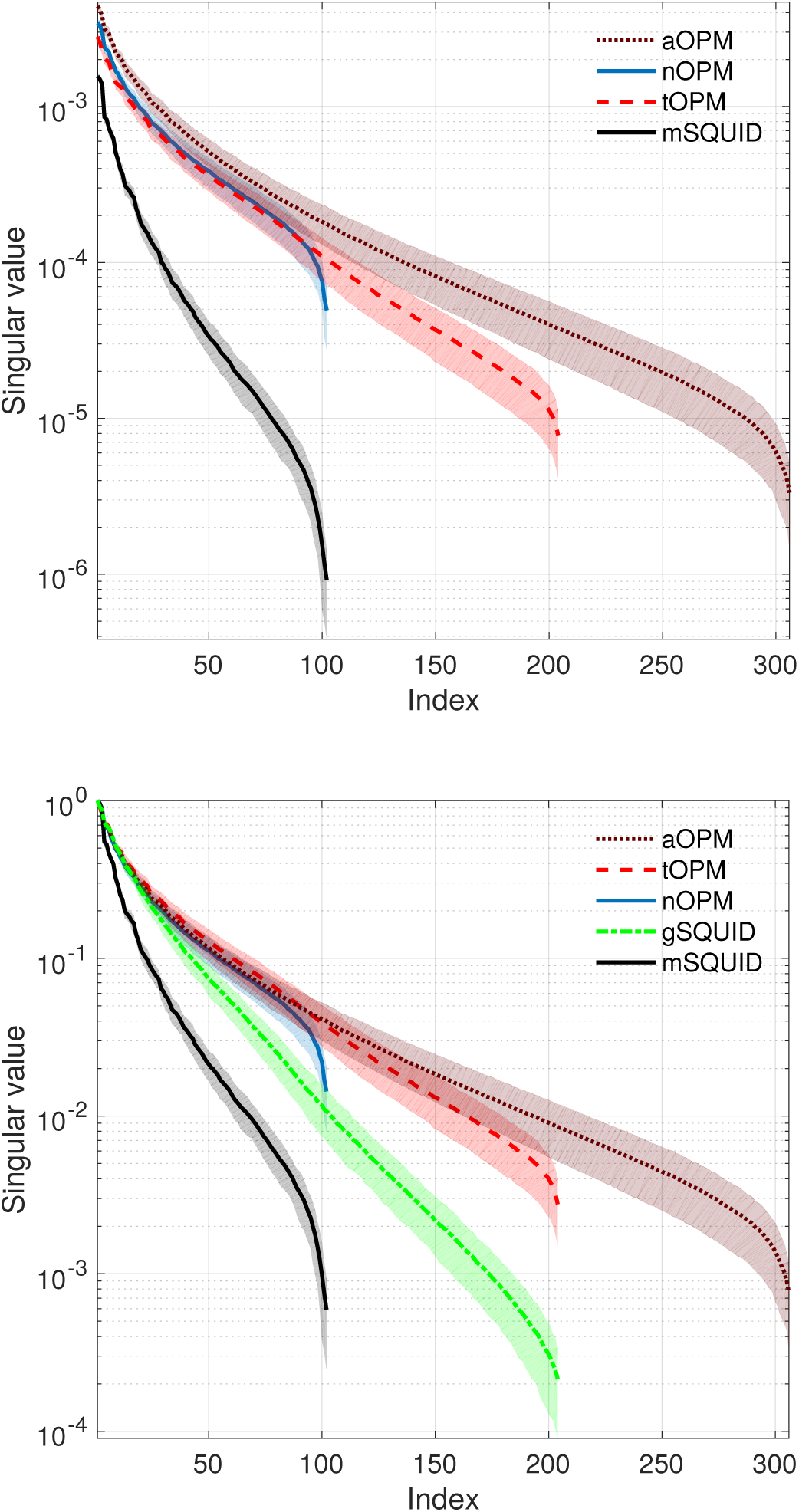
Singular-value spectra of the lead-field matrices (top) without normalization and (bottom) normalized by the maximum singular value of each array. Singular values were averaged across ten subjects; the shaded regions give ±1 standard deviations.

